# Emerging viruses in British Columbia salmon discovered via a viral immune response biomarker panel and metatranscriptomic sequencing

**DOI:** 10.1101/2020.02.13.948026

**Authors:** Gideon J. Mordecai, Emiliano Di Cicco, Oliver P. Günther, Angela D. Schulze, Karia H. Kaukinen, Shaorong Li, Amy Tabata, Tobi J. Ming, Hugh W. Ferguson, Curtis A. Suttle, Kristina M. Miller

## Abstract

The emergence of infectious agents poses a continual economic and environmental challenge to aquaculture production, yet the diversity, abundance and epidemiology of aquatic viruses are poorly characterised. In this study, we applied salmon host transcriptional biomarkers to identify and select fish in a viral disease state but only those that we also showed to be negative for established viruses. This was followed by metatranscriptomic sequencing to determine the viromes of dead and dying farmed Atlantic (*Salmo salar*) and Chinook (*Oncorhynchus tshawytscha*) salmon in British Columbia. We found that the application of the biomarker panel increased the probability of discovering viruses in aquaculture populations. We discovered viruses that have not previously been characterized in British Columbian Atlantic salmon farms. To determine the epidemiology of the newly discovered or emerging viruses we conducted high-throughput RT-PCR to reveal their prevalence in British Columbia (BC), and detected some of the viruses we first discovered in farmed Atlantic salmon in Chinook and sockeye salmon, suggesting a broad host range. Finally, we applied *in-situ* hybridisation to confirm infection and explore the tissue tropism of each virus.

## Introduction

The number of documented RNA viruses is undergoing rapid growth, largely since the next generation sequencing and associated metatranscriptomic boom of the last decade (1–5). As a result, we are gradually uncovering previously unknown viral diversity, known as viral dark matter (6, 7). However, metagenomics has limitations, and there are barriers to viral discovery – screening a large number of samples is costly and time consuming and viral genomes are often outnumbered by orders of magnitude amongst the host, bacteria and other contaminants (8–10). Furthermore, huge amounts of data are generated in a ‘sequence and see’ approach to metagenomics which can be costly and time consuming to analyse, and there is no guarantee that the genomes that are found are biologically relevant, or even infect the host which was sampled.

Traditionally, the most common diagnosis of fish viruses uses culture of virus from diseased host material in a monolayer cell line – a method suited to screening of large numbers of diseased tissues (high throughput). Cultures are monitored for the development of cytopathic effects (CPE), this being the first step in the identification of the virus. The majority of documented fish viruses produce obvious CPE, but most were discovered via culturing, and this selection bias means there is the strong potential to miss novel virus groups that are ***not*** amenable to culture (6). Meanwhile, our knowledge of viruses in nonmammalian vertebrates is comparatively scarce, but growing rapidly, and most families of RNA viruses once thought to be restricted to mammals are now known to infect or be associated with fish metatranscriptomes, some of which appear to be ancient relatives of important human pathogens (1, 11, 12).

The global demand for seafood is rising, whilst many wild fisheries are decreasing in productivity (13). There is extensive interest in the potential impact of infectious disease on wild salmon populations. The 2009 decline of Canada’s largest salmon fishery, Fraser River sockeye salmon, instigated a public enquiry into declining salmon stocks, in which infectious disease was highlighted as an area deserving of more study (14). Meanwhile, a recent federal report found that half of Canada’s Chinook populations are endangered, with nearly all other populations in decline (15). Aquaculture production is growing rapidly to meet the increased seafood demand, and its expansion provides increased opportunity for the transmission of emerging viruses and the evolution of virulence (16, 17). Disease can be a major factor limiting aquaculture production (18) and lead to substantial economic loss (19, 20). Although there is a growing body of research on the risk posed by pathogens and parasites associated with aquaculture on wild marine species (21–25), there has been little research that considers the risk of emerging viruses in this context. In fact, a recent government audit criticised government regulatory bodies for not addressing this knowledge gap and the potential risk to wild salmon from emerging viruses associated with the aquaculture industry (26).

Factors affecting salmon populations are multi-faceted and complex, but accumulating evidence suggests that infectious disease may play a role in the collapse of wild salmon populations in the Eastern Pacific. A range of pathogens have been linked to mortality in adult wild Pacific salmon (23, 24, 27), but there are few data on the role of infectious diseases in the >90% mortality of migratory juvenile salmon in the ocean. Gene expression profiles consistent with an immune response to viruses have been associated with mortality in wild migratory smolts and adults (28, 29), as well as in unexplained aquaculture mortalities of salmon in marine net pens in BC (30, 31). Together, these suggest there are a range of viruses that may contribute to decreased survival of migratory salmon in BC. Moreover, there are concerns about the impact of emerging infectious agents associated with the expansion of salmon enhancement hatcheries (12, 32) and salmon aquaculture in BC, which operate in the same waters through which wild Pacific salmon migrate (33).

Studying disease in wild populations is exceedingly complex (34), and although terrestrial strategies can include treating for known infectious agents, fish health investigations often begin with discovery. In the ocean, mortality events are rarely observed; our sampling efforts solely capture live fish, and weak and dying fish are probably predated before the disease progresses to mortality (27, 30). In order to surmount some of these difficulties, and to select samples for virus discovery, we applied a viral disease development (VDD) biomarker panel shown to be predictive of an active RNA virus infection in salmon (30) followed by metatranscriptomic sequencing, molecular surveillance and *in-situ* hybridisation to prove infection.

## Results and Discussion

### Viral disease development (VDD) biomarkers and virus discovery

The VDD panel of host transcriptional biomarkers is capable of differentiating fish in an active RNA viral disease state from those carrying bacterial or fungal disease (30). For sequencing, we selected farmed Atlantic and farmed Chinook salmon which exhibited a strong viral disease development (VDD) signal, but were negative for any RNA viruses known to infect salmon in the North Pacific. For example, a previous study showed that one third of a population of moribund Atlantic salmon were found to be in a viral disease state (31), and as half of these fish were negative for RNA viruses we assumed that they contained undiscovered RNA viruses. Metatranscriptomic sequencing was carried out on dead and dying farmed Chinook and Atlantic salmon collected as part of the Canadian Department of Fisheries and Oceans aquaculture audit, which samples marine finish aquaculture facilities to meet regulatory requirements. Illumina sequencing, followed by de novo assembly and a similarity search (translated BLAST to the nr database) revealed a collection of novel viruses or novel variants of existing viral species, including the first detection of Atlantic Salmon Calicivirus (ASCV) in North America, a new strain of Cutthroat trout virus (CTV-2), and three putative RNA viruses (pNarnaV, pTotiV, pRNAV).

To demonstrate the effectiveness of sample selection via the VDD biomarker, we calculated the probability of detecting each of the novel viruses if we had chosen a random subset of the aquaculture audit samples available for sequencing. In this analysis, we also included the recently discovered Salmon pescarenavirus 1 and 2 (SPAV), Chinook Aquareovirus (CAV) and Pacific salmon nidovirus (PsNV) (12). The probability of detecting a novel virus depends on how many samples were sequenced and how common the virus is in the dataset (Figure 1a). The frequency is based on NGS-detectable infections, i.e. if 1,000 fish were sequenced and 100 were positive for a specific virus, the prevalence would be defined as 10%. Similarly, figures 1B and 1C show curves for the probability that randomly sequencing k samples would lead to virus detection in at least one salmon, but in these plots, observed detections by RT-PCR in the aquaculture data sets were used to define prevalence. Only viruses that had positive detections in the aquaculture audit for each species were included. A theoretical copy number threshold of 1,000 nucleic acid copies per μg of tissue was used to estimate the chance of identifying a novel virus in a metatranscriptomic sequencing run. With this threshold, five of the viruses were detected in the Chinook salmon audit data (9/203 with SPAV-1, 7/203 with PsNV, 3/203 with CTV-2, 2/203 with pNarnaV and 24/203 with CAV) and three of the viruses were detected in the Atlantic salmon audit data (439/665 with ASCV, 320/ 665 with CTV-2 and 9/665 with pNarnaV). Both ASCV in Chinook salmon and SPAV-1 and CAV in Atlantic salmon had positive detections but were below the copy number threshold of 1,000; thus the corresponding curves show probabilities of zero. pTotiV (described below) was first discovered in an alternative aquaculture data-set (31) which was not part of the audit, so is excluded from this analysis. The probabilities described in Figure 1 demonstrate that for more widely occurring viruses, a targeted sequencing approach is not necessary; whereas, the rarer viruses would likely remain undiscovered without our VDD-based sample selection approach.

**Figure 1.**
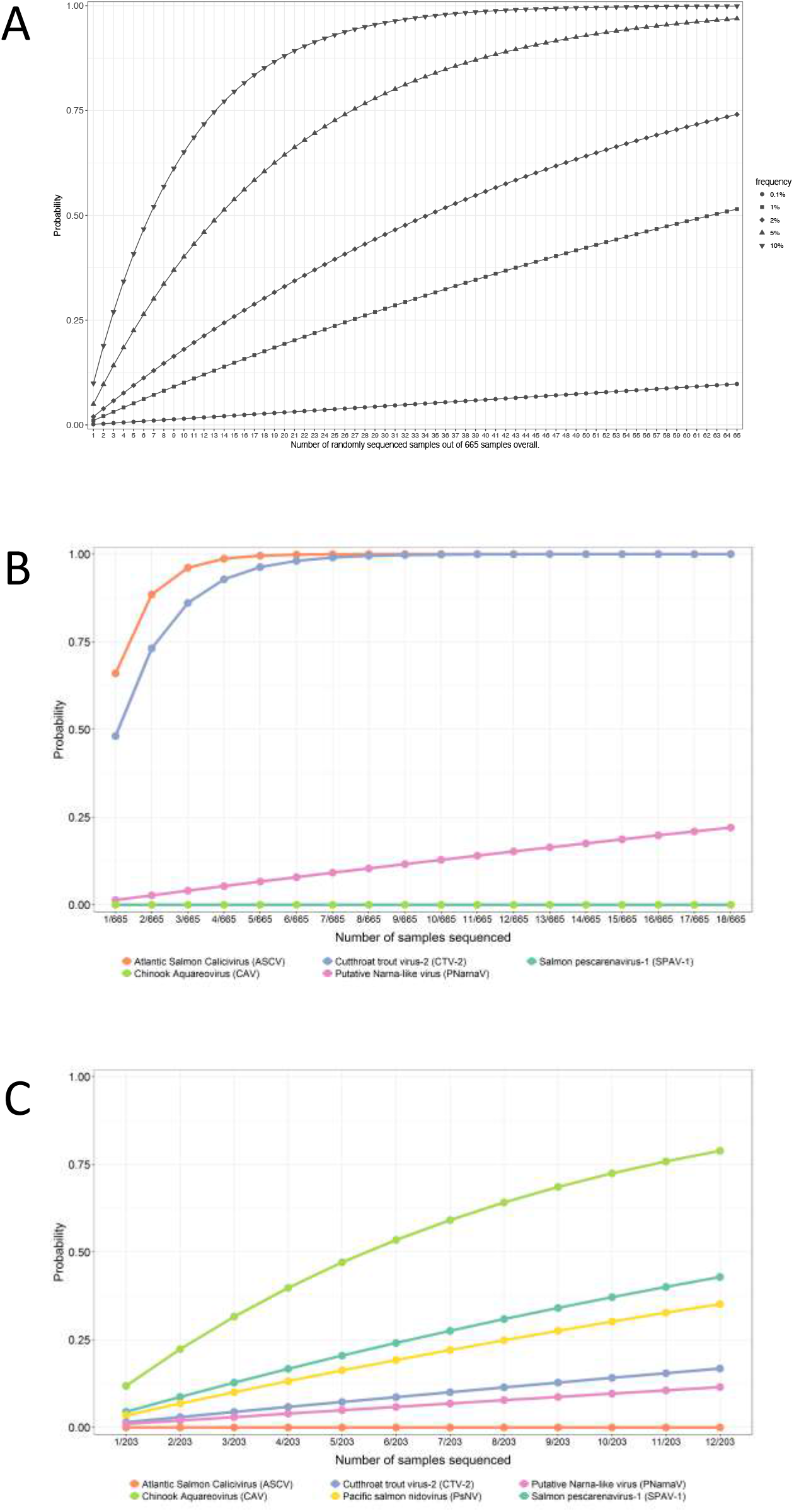
**A)** Probability of detecting a novel virus in 665 samples by randomly sequencing the specified subset of samples. Curves represent different scenarios where the specified percentage of 665 samples overall has a theoretical detectable viral load for NGS sequencing. **B)** Probability of finding the specified virus when sampling k samples randomly from the set of 665 Atlantic salmon in the farm audit data and **C)** when sampling k samples randomly from the set of 203 Chinook salmon in the farm audit data. A copy number threshold of 1,000 was used to define detection. The x-axis goes up to values of k=12 for Chinook salmon and k=18 for Atlantic salmon, which were the number of samples sequenced based on suggestions derived from VDD patterns in the audit data sets. Of these, we did not detect a novel virus in 2/18 Atlantic salmon and in 5/12 Chinook.

The number of viral transcripts in fish metatranscriptomes is hugely outnumbered by host transcripts and other contaminants (10, 35). Therefore to achieve full genome coverage, high depth of sequencing is required. In viral discovery studies, there is a trade-off between sequencing a larger number of samples at a lower depth of coverage versus less samples at a higher depth. If coverage is too low, viral genome assembly can be fragmented and only regions of the genome with similarity to known viruses are detected whilst others could be missed entirely. As a compromise, our strategy was to sequence a larger number of VDD positive samples at a relatively lower depth. Although this did not yield ‘coding-complete’ sequences for all the putative novel viruses, we were able to sequence enough of the viral genome to design RT-PCR primers to inform a high throughput epidemiological survey in which we screened over 9000 salmon sampled over one decade along the coast of Southern BC (figure 2), conduct phylogenetic analyses (figure 3), as well as design probes for in situ hybridisation to determine infectivity and tissue tropism (figure 4).

**Figure 2.**
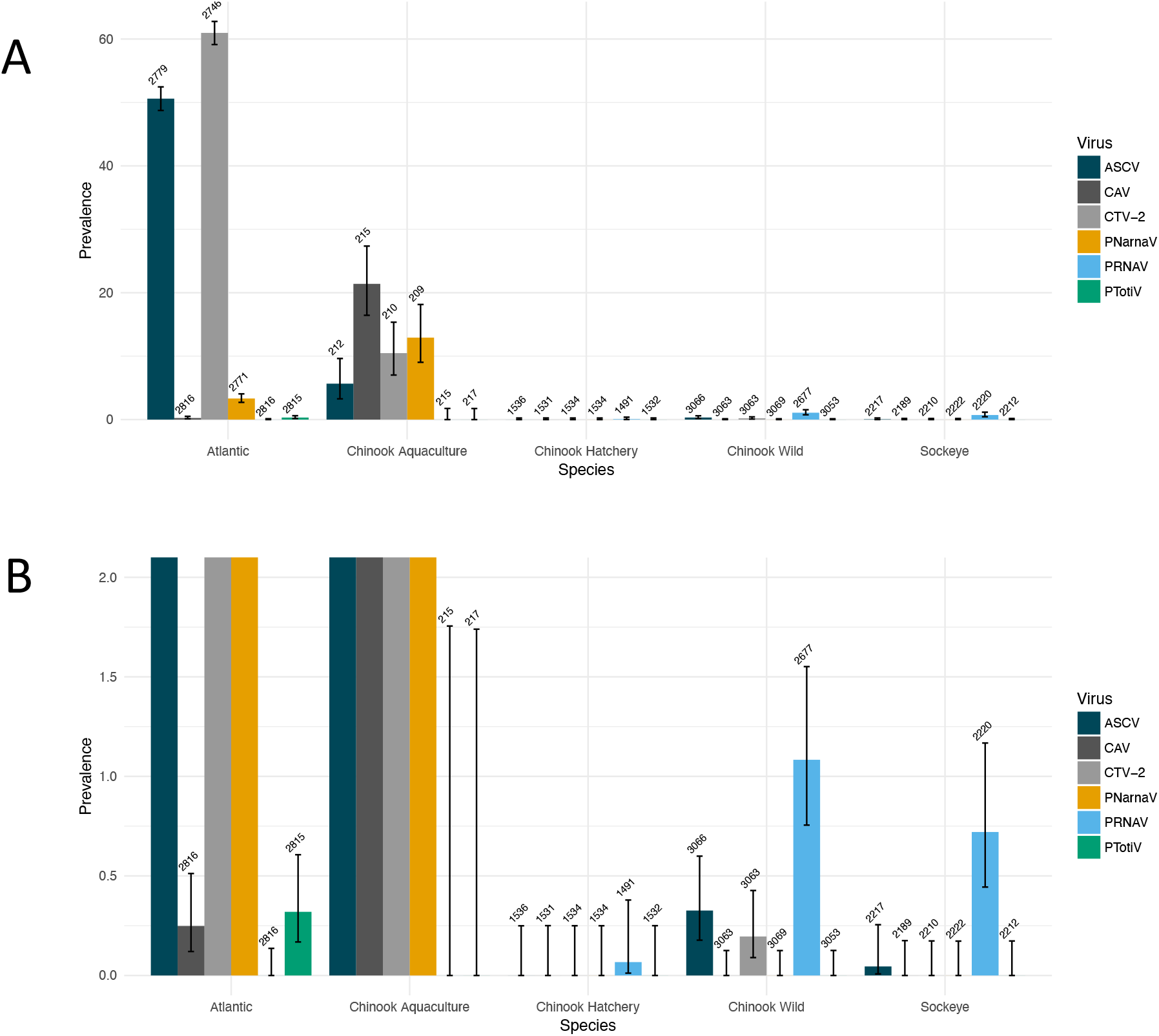
**A)** Prevalence (%) of the emerging viruses detected by high-thoughput RT-PCR. Numbers show the total number of samples tested, and the error bars show the confidence intervals on the calculated prevalence using the Wilson method. **B)** displays the same data but the prevalence (y-axis) is expanded (maximum prevalence of 2%) to show the detection of rarer viruses.

**Figure 3.**
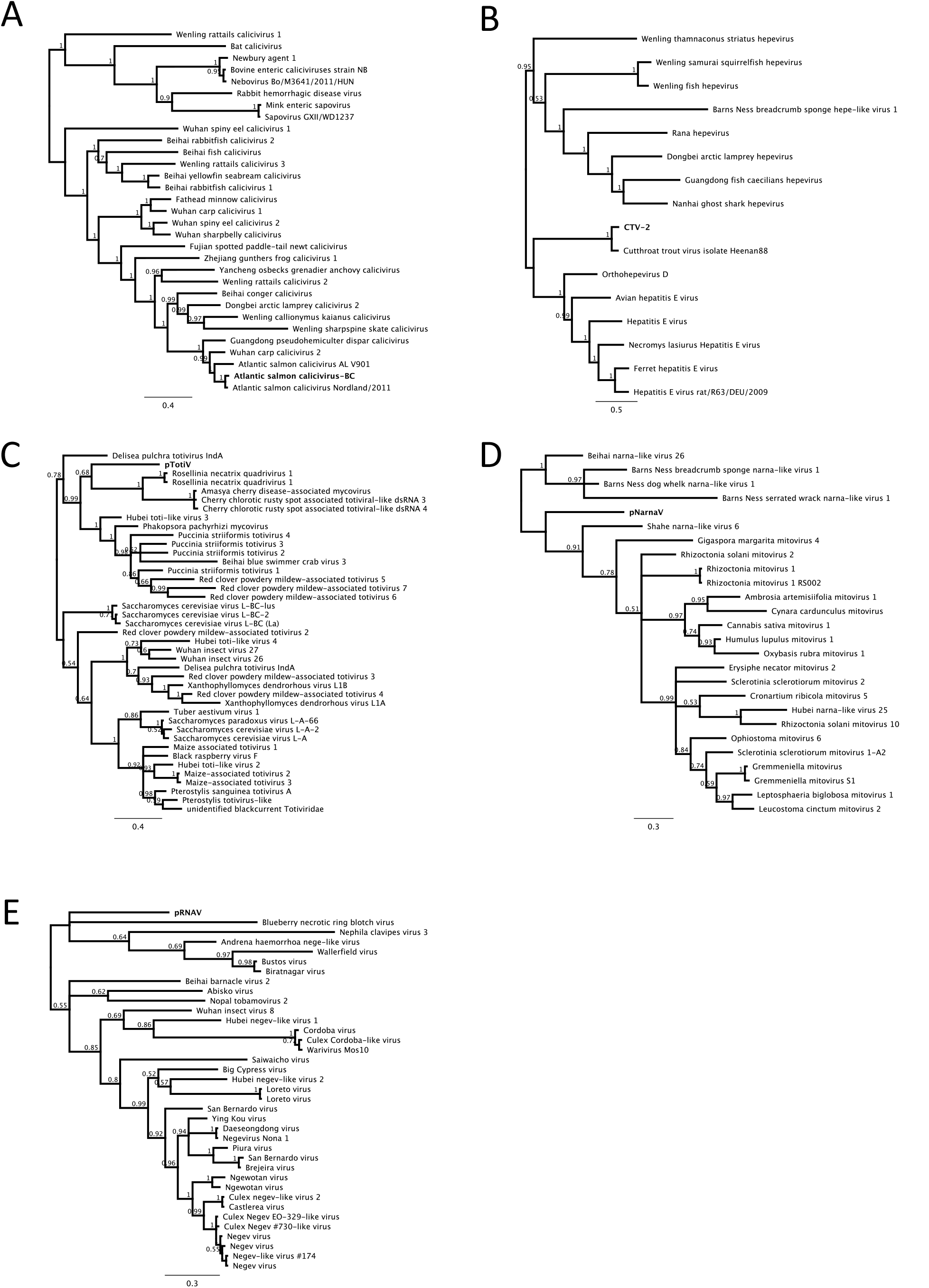
Phylogenetic relationship of **A)** Atlantic salmon calicivirus **B)** Cutthroat trout virus −2 (BC) strain **C)** putatative totivirus (pTotiV) **D)** putatative narna-like virus (pNarnaV) and **E)** putative RNA virus (pRNAV). Sequences from this study are shown in bold. Phylogenies are based on the predicted amino acid RdRp encoding sequence. Branch labels show the posterior probability as calculated by MrBayes and the tree is mid-point rooted for clarity only.

**Figure 4.**
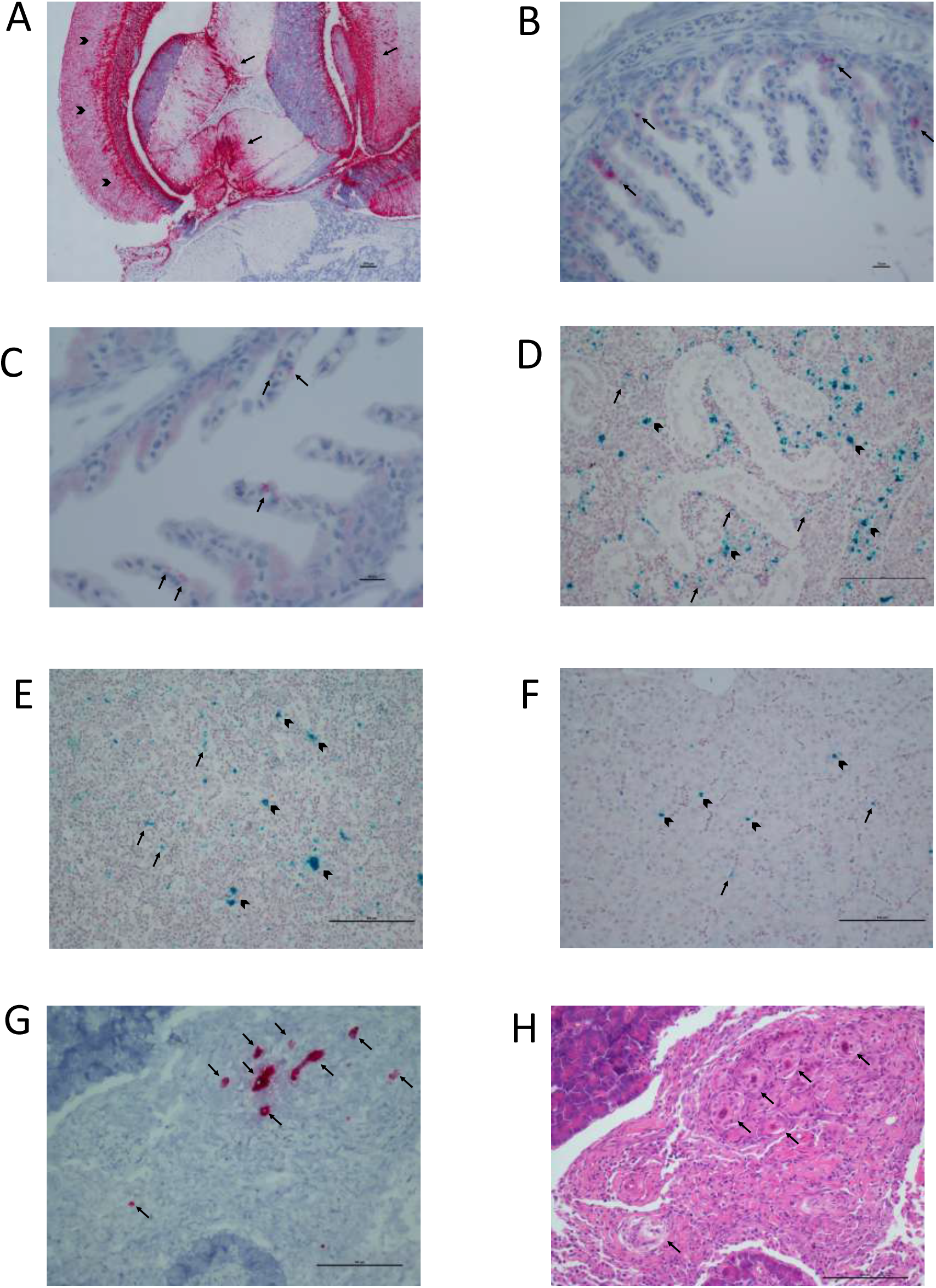
**A)** CTV-2 in Atlantic salmon: intense staining (red) localizing viral particles in the optic lobes, particularly the stratum griseum, periventriculare and fibrosum profundum (arrowheads) and most of the mesencephalon (arrows). Scale bar 100μm. **B and C)** PsNV in Chinook salmon: viral particles (red, arrows) localized in the epithelial cells of the gills. Scale bar 10μm. **D)** CAV in Chinook salmon: positive marking observed on macrophages (arrowheads) and other cell types in the erythropoietic tissue (arrows). Scale bar 100μm. **E)** CAV in Chinook salmon: widespread marking (green) in the spleen (arrows), particularly in the macrophages (arrowheads). Scale bar 100μm. **F)** CAV in Chinook salmon: positive marking (green) observed in red blood cells (arrows) but also inside some hepatocytes (arrowheads). Scale bar 100μm. **G)** PRNAV-1 in Chinook salmon: serosal surface of intestine showing granulomatous inflammatory response engulfing embryonated metazoan eggs (probably trematodes), marked (red, arrows) for viral particles. Scale bar 100μm. **H)** PRNAV-1 in Chinook salmon: same section as G), showing granulomatous inflammation (arrows) encompassing several embryonated metazoan eggs (probably trematodes) infected with virus. Scale bar 100μm

### A*tlantic Salmon Calicivirus (ASCV)*

ASCV was the first calicivirus in fish to be fully characterised and sequenced (36). ASCV was detected in a high proportion of farmed Atlantic salmon in Norway, and was observed in association with several diseases including heart and skeletal muscle inflammation (HSMI). However, ASCV was also commonly found in healthy fish and a more recent study found no correlation between ASCV and HSMI (36, 37). However, there is evidence that co-infection of caliciviruses with other viruses could be linked to clinical manifestations of disease in baitfish (38). In this study, in farmed Atlantic salmon from BC, we discovered viral genomes extremely closely related to the ASCV genome first discovered and sequenced in Norway (36). Despite the high prevalence of ASCV in farmed fish in Norway, it was only recently reported in Atlantic Canada (Teffer et al. *in press*), and this represents the first published detection of ASCV in the Pacific at the time of writing, which is surprising considering it was found to infect over half of the Atlantic salmon we tested (figure 2).

Caliciviruses are single-stranded, positive-sense RNA viruses which cause a wide variety of diseases in mammals and birds (39–43), and were initially thought to result in transient clinical disease such as blistering of the skin, pneumonia, abortion, encephalitis, myocarditis, hepatitis, and diarrhea, but have now been associated with hemorrhaging and mortality (44–46). Phylogenetically, caliciviruses are part of the picorna-like virus group which is thought to have evolved since eukaryogenesis (47), so it is not surprising that they are found extensively in all groups of vertebrates. Interestingly, some caliciviruses infect marine mammals (48), and remarkably, at least one fish species has been reported as reservoir for vesicular exanthema of swine virus (VESV), which can infect and cause disease in mammals (49, 50). More recently there has been a huge surge in calicivirus discovery in fish and amphibia (1, 36, 38). Phylogenetically, fish caliciviruses are basal to the much better studied mammalian caliciviruses (1). As expected, ASCV groups within a distinct clade from the mammalian caliciviruses (Figure 3A), and is more closely related to metatranscriptomic discovered fish caliciviruses (1) and fathead minnow calicivirus (38), a novel calicivirus isolated from farmed baitfish in Wisconsin, USA. It is becoming clear that caliciviruses are widespread in fish, and although we know very little of their direct role in lesion development, fish caliciviruses have been associated with diseased fish in an aquaculture setting (38). These newly described fish caliciviruses (1, 38) are phylogenetically distinct from those first described in mammals, suggesting that spillover from fish to mammals is not a common occurrence, as has been previously suggested (49, 50).

The initial discovery of ASCV (36) revealed two variants that shared 70.9% sequence identity, one of which was sequenced directly from field material while the other was isolated through cell culture and subsequently sequenced. In this study, the coding-complete genome sequence of ASCV (referred to as ASCV-BC1) was assembled from a sample taken from a farmed Atlantic salmon (G651). Interestingly, the variant sequenced in farmed Atlantic salmon from BC is more closely related to the Norwegian field variant than the cell-culture isolate (figure 3A). These viruses share 83.8% nucleotide identity and 95.7% amino-acid identity (in ORF1), and form a distinct clade from the cell-culture isolate based on their aminoacid sequence. This suggests that the cell culture isolate may have undergone selection pressure under culture conditions, which is plausible considering it was cultured in the GF-1 cell line, which originates from grouper, *Epinephelus coioides* (51). As expected for two closely related viral genomes, the ASCV-BC1 variant we sequenced shares a genome structure similar to those in Norway, with one large open reading frame (ORF 1; 7086 nt), and a second, smaller open reading frame (ORF 2; 378 nt) which overlaps with ORF1 in the 3’ end of the genome (figure S1). ORF2 of ASCV-BC1 is highly conserved and shares 98.4% amino acid identity with the Norwegian field strain of ASCV. Due to the sequence similarity of this sequence to the Norwegian variant, and its high prevalence in farmed Atlantic salmon in BC, it seems plausible that this virus originates from Norwegian Atlantic salmon that were introduced to BC for aquaculture.

Infectivity of ASCV for Atlantic salmon has previously been shown in Norway (36), but its presence in BC was until now unexplored. Surveillance by high-throughput RT-PCR found that ASCV was common in dead and dying farmed Atlantic salmon (1406 of 2779 fish). Interestingly, ASCV was detected in 12 of 212 farmed Chinook and in 10 of 3066 wild Chinook (figure 2), suggesting a broad host range and corroborated by replication of ASCV in a grouper cell line (36). Transmission between populations and species is entirely feasible considering caliciviruses remain viable even after exposure to sea water for 14 days.

### Cutthroat trout virus-2 (CTV-2)

In the past decade, there has been a surge in discovery of Hepatitis E viruses in various animals (52), including Cutthroat trout virus (CTV), the first hepevirus identified in fish (53–55). CTV was first isolated in the CHSE-214 (Chinook salmon embryo) cell line, from material collected from cutthroat trout (*Oncorhynchus clarkia*) (55), and it is the first member of a proposed new genus *Piscihepevirus* (56, 57). To date, CTV has not been associated with disease but it is widespread in trout populations in the Western USA; it has also been detected in Atlantic salmon (55, 58). It has been suggested that CTV should be renamed Piscihepevirus-A, as it is capable of infecting species other than cutthroat trout (56). More recently, a range of hepeviruses have been discovered in fish metatranscriptomes. These viruses are phylogenetically diverse, and are likely representative of other hepevirus genera which infect fish, but these remain unclassified. Here we discovered a viral sequence bearing similarity to CTV which we named CTV-2 due to a relatively high sequence similarity and overlapping host range.

Due to rapid replication and high error rates, RNA viruses exhibit vast diversity (59), resulting in genetically highly heterogeneous population structures referred to as viral quasispecies. The host range of a virus can be affected by the level of diversity within a quasispecies (60). Rather than a randomly generated swarm of mutants around one variant, quasispecies commonly consist of a number of master variants, each with their own swarm of random mutations (61, 62). These high levels of heterogeneity enable RNA viruses to occupy large areas of sequence space and likely explains the broad host range of CTV (55, 59). Figure 3B shows the phylogenetic placement of CTV-2 within the wider *Hepeviridae* family and in relation to the recent hepeviruses in fish and frogs identified from metatranscriptomic data (1). The genetic diversity between CTV and CTV-2 is relatively small in the wider context of the diversity within the *Hepeviridae*. We hypothesise that CTV-2 represents a second master variant of the CTV quasispecies adapted to salmonid species. Although many sequence variants of CTV have been partially sequenced (55), CTV-2 represents a new master variant, which is phylogenetically distinct from all published CTV variants (supplementary figure 1). CTV-2 shared 73.7% nucleotide identity (82% amino-acid) to the published full-length CTV genome, whereas previous studies found that other variants of CTV had a nucleotide diversity of less than 8.4% (55).

Notably, CTV-2 is missing a homolog of open reading frame 3 (ORF3) (supplementary figure 2) which is possessed by the original CTV sequence (55). Predicted ORFs are fragmented in the region where ORF3 is expected. In other hepeviruses, ORF3 encodes an immunogenic protein which is thought to be involved in virion morphogenesis and viral pathogenesis, but its role is not fully understood. ORF3 is not highly conserved among hepeviruses and there is a low similarity in ORF3 encoding sequences between viruses from different hosts, suggesting it represents viral adaptation to a particular host (54). Furthermore, hepeviruses recently sequenced in fish are also missing ORF3 (1).

*In-situ* hybridisation revealed that CTV-2 infection was systemic, although preferential sites appeared to be the optic lobes of the brain (particularly the stratum griseum periventricular and the stratum fibrosum profundum) and the whole mesencephalon area (Figure 4A). The same areas were associated with mild neuronal necrosis and degeneration as well as mild ventriculitis and proliferation of ependymal cells. The virus was also observed in the cardiomyocytes (some of which were necrotic), in the splenic ellipsoids and in the renal hemopoietic tissue.

CTV-2 was prevalent in the salmon aquaculture samples, and was detected in 60.1% of farmed Atlantic salmon and 10.5% of farmed Chinook salmon (figure 2). However, samples with a high viral load were found in Atlantic salmon only, and these were mostly fish with a strong viral disease biomarker signal. Similar to ASCV, detection of this virus in Chinook highlights the potential for cross-species transmission. Interestingly, CTV is widespread in North America, but has not been detected on any other continent (55), which suggests this may represent a transmission event from a Pacific species to Atlantic. However, a closely related virus has been detected in Atlantic Canada (58), so the origin is not entirely certain.

### Pacific salmon nidovirus (PsNV)

PsNV, first described in a metatranscriptomic survey of Pacific salmon (12), was shown to be predominantly observed at high prevalence over multiple years in Chinook salmon leaving freshwater hatcheries, and was localised to the gills by RT-PCR. Here, *in-situ* hybridisation was used to localise the viral RNA to the epithelial cells of the secondary lamellae of the gills, some of which showed swelling and hydropic degeneration (Fig. 4B & C). This finding supports the hypothesis that disease caused by the virus is manifested when fish undergo physical stress due to osmotic imbalance as they move from fresh to marine waters during smoltification (63).

### Chinook Aquareovirus (CAV)

We previously used metatranscriptomic sequencing of aquaculture Chinook to reveal a novel reovirus, Chinook Aquareovirus (CAV) (12); phylogenetic analysis of the RdRp predicted that CAV is part of the genus *Aquareovirus*. In this study, *in-situ* hybridisation shows that the virus had a systemic distribution, although it was primarily observed in the splenic, peri-sinusoidal macrophages and in the renal portal macrophages (Figure 4D, E & F). It is unclear if CAV infects these immune cells as a ‘Trojan horse’, and mediates virus spread in the whole organism while being protected from the host immune response (64), or whether the virus was simply localized in macrophages following phagocytosis in an effort to eliminate the virus from the blood. Additionally, scattered hepatocytes showed viral particles in the cytoplasm, although lesions were not visible in these cells.

CAV can be grouped with a growing number of aquareoviruses, which are phylogenetically basal to those first discovered in fish (12, 65). This group includes Hubei grass carp reovirus (HGCRV, also known as GCRV104) (66) and Pangasius aquareovirus (unpublished, sequence available on GenBank) and the Western African Lungfish Reovirus (1). Of these, HGCRV is the most studied, as it causes hemorrhagic diseases of grass carp and serious loss to carp aquaculture in China (67). Interestingly, the addition of these metatranscriptomic sequences (1, 66) further breaks down the shallow distinction between aquareoviruses and orthoreoviruses, lending weight to the call for these genera to be merged (65). CAV is more akin to the aquareovirus group due to its phylogenetic placement based on the RNA dependent RNA polymerase (RdRp) as well as possessing 11 genome segments, each of which appear to encode an open reading frame (ORF). The segments were named in accordance with their homology to the HGCRV segments (supplementary table 1). All segments are coding complete but lack the terminal conserved untranslated region typical of reoviruses (68). Nine of the 11 genome segments were identified by their homology to other aquareoviruses and the presence of conserved reovirus domains (supplementary table 1), but there were no hits to the outer fiber or FAST proteins found in other aquareoviruses and orthoreoviruses.

As segments 7 and 11 showed no recognizable sequence similarity using a translated BLAST of the RefSeq non-redundant database (69), these sequences were attributed to CAV by their size, predicted ORFs, and their primary protein structure. Further analysis of these predicted proteins revealed features reminiscent of closely related reoviruses. For example, HHPRED analysis of segment 7 revealed a predicted protein domain similar to the orthoreovirus sigma-1 protein and adenovirus fiber proteins, suggesting it encodes the fiber protein that is present in some, but not all aquareoviruses and Piscine orthoreovirus (65). Similarly, the putative segment 11 of CAV, contains a signal peptide and transmembrane domains as detected by Phobius (70) (supplementary figure 3, supplementary table 2). These observations are similar to those of HGCRV, Piscine orthoreovirus and GCRV-HZ08/GD10, which also contain transmembrane domains that could represent a membrane interacting non-structural protein (65).

### Other RNA viruses

Finally, smaller fragmented assemblies showed homology to other groups of RNA viruses were sequenced from three different samples. Although there was not full genome coverage, we were able to design RT-PCR assays for each putative virus to see how widely distributed they were in farmed and wild salmon.

For all three samples, the assay was designed on a *de novo* assembled sequence that showed similarity to the RdRp, enabling phylogenetic analysis. Although these RdRp sequence fragments were under 500 nt, they are sufficient to reveal phylogenetic relationships of viruses in fish (35). Each assembly was translated and aligned with protein sequences obtained from GenBank using search results from BLAST to deduce the phylogenetic relationships of these putative viruses.

On the whole, these viral transcripts were more closely related to invertebrate-associated groups of viruses, implying they were more likely to be infecting invertebrates associated with the fish, rather than the salmon themselves, as discussed in (35). For example, several small assembled sequences, referred to in this paper as PRNAV-1 (putative RNA virus-1) (Accession numbers MN995809-MN995814) were detected in a wild Chinook smolt (Sample B2175). These sequences were identified as viral as they showed similarity to a group of arthropod-associated viruses in the proposed *Negevirus* taxa (figure 3E) and the arthropod *Iflavirus* genus. Additionally, screening by high-throughput RT-PCR revealed that this virus was detected in only 0.42% of Chinook and sockeye salmon (57 of 13296 samples) (figure 2). Interestingly, recent evidence of arthropod viruses circulating in the blood of a vertebrate host (71) suggests that crosskingdom transmission could be more common than previously thought. Similarly, there have been reports of novel flaviviruses infecting both marine vertebrates and invertebrates (72). Here we show, using *in-situ* hybridisation, that this putative virus is not an environmental contaminant, but appears to be associated with an invertebrate parasite of the salmonid host, raising the possibility that there is an alternative invertebrate host for this virus. The virus was primarily observed in structures (tentatively identified as embryonated metazoan eggs, possibly trematode eggs) encysted in the lamina propria of the intestine and localised on the serosal surface. The putative eggs were surrounded by a granulomatous inflammatory reaction. Viral RNA was also detected in the exocrine pancreas and peri-capsular adipose tissue of the spleen. It is not clear if the localisation reflects the dissemination of the virus from the metazoan eggs in the intestine, or infection of an immature stage of the parasite itself. The detection in fish of viruses that phylogenetically resemble viruses of invertebrates reflects recent work which found the sequence from the same virus in both crabs and sharks (72). It is apparent that in the marine environment, certain groups of viruses are transmitted horizontally between invertebrates and vertebrates, in which case distinction between groups of viruses that infect vertebrates and invertebrates is starting to break down.

Additionally, we identified sequences with similarity to viruses belonging to the *Narnaviridae*, a family of viruses that lack a capsid, which we named PNarnaV (putative Narna-like virus), due to its phylogenetic placement (Figure 3D). This sequence was detected solely in aquaculture samples, both Atlantic (3.3% prevalence) and Chinook (13% prevalence). Narnaviruses are non-infectious RNA transcripts that are assumed to be vertically transmitted and which reproduce inside the mitochondria (genus *Mitovirus*) or in the cytosol (genus *Narnavirus*) of the host (73, 74). These viruses are known to infect fungi, and are widespread in the viromes of plant-pathogenic fungi (75). This group was greatly expanded in a recent metatranscriptomic study of invertebrates (2), and many of these viruses appear to originate from host-associated fungi, although it has been suggested that the sheer number and diversity of this group of viruses implies a broader host range (73). It is unclear if the putative narna-like virus is infecting the salmon, or an associated fungal parasite, but as the sequence was only found in dead salmon, it suggests the virus may be associated with fungi only associated with decomposing fish.

Similarly, a small sequence assembly (309 nt) with similarity to a variety of fungi-infecting totiviruses, was detected in farmed Atlantic salmon and named PTotiV (putative toti-like virus) (figure 3C). Originally, totiviruses were thought to only infect unicellular fungi, but the known host range of totiviruses has expanded to include arthropods and fish (76, 77). Pertinently, a totivirus, piscine myocarditis virus (PMCV), was shown by *in-situ* hybridisation to infect Atlantic salmon and is associated with cardiomyopathy syndrome (77, 78). Additionally, a totivirus has been implicated as an emerging pathogen in paenid-shrimp aquaculture (79, 80). As there was little to no recognizable sequence similarity between pTotiV and PMCV it implies that pTotiV is evolutionarily distinct from PMCV, and may represent a second evolutionary lineage of totiviruses infecting vertebrates. However, the prevalence of pTotiV was extremely low, with just 9 out of 2815 samples of farmed Atlantic salmon samples positive for the virus, and no positive detections in 4802 samples of Chinook salmon. There is a developing body of work demonstrating that within one viral family, cross-Kingdom host ranges can occur (81, 82). These findings highlight the difficulties in identifying the host of novel viruses discovered in metatranscriptomic surveys of RNA viruses purely based on sequence alone.

### Conclusions

In this study, we used *in situ* hybridisation, sequencing and RT-PCR surveillance to show that ASCV-BC1 and CTV-2 occur in salmon, and have phylogenetic proximity to other viruses that infect fish, suggesting that they may be important to salmon health. However, the detection of RNA does not demonstrate infection, and in some cases novel virus sequences could represent a virus from an alternative host associated with salmon, such as PRNAV-1, which was seen in embryonated eggs of what was likely a trematode parasite. Similarly, the hosts of the putative viruses, pTotiV and pNarnaV, are not clear, and they may have no relevance to investigations of salmon health.

We show that the viral disease development (VDD) biomarker panel increased the probability of discovering previously unknown viruses. In the future, the VDD panel will be an important tool to enable the discovery of viruses in wild salmon, in which we expect to find fewer positive or high-load samples than in farmed fish. Moreover, given the high overlap in genes contributing to the salmon VDD panel and those discovered to be predictive of respiratory viral diseases in humans (83, 84), there is a high potential that a similar panel of genes could be applied for viral discovery in a diverse range of vertebrate host species (30).

Disease is a factor affecting rates of population mortality, and infectious diseases are an integral part of ecosystems. However, the presence of a virus is not synonymous with disease; infections can be acute, subacute, chronic or inapparent (85). However, changing environmental or anthropogenic factors, for instance, the introduction of a vector (86), an alternative host or high density rearing environments (17) can facilitate rapid viral propagation and result in increased evolutionary pressure for the emergence of more virulent strains or enable host-range expansion. Therefore, to understand the mechanisms of viral emergence, we need to characterise virus diversity amongst different host species. Moreover, environmental stressors such as climate change, could result in disease emergence (87, 88), even from agents that might have previously been tolerated. Therefore, continual discovery and surveillance of emerging viruses in these ecologically important and fragile species will be vital for management of both aquaculture and wild resources in the future.

## Materials and Methods

### Nucleic acid extractions

We applied similar sequencing and screening approaches to those used in a previous study (12), but details are included below for completeness. Samples were provided by the Fisheries and Oceans, Canada Aquaculture Management Division, Environmental Watch Program, High Seas Program, Strait of Georgia salmon Program, PARR Program and Salmon Enhancement Program as well as by the Hakai Institute. Hatchery samples are identified by fin clipping, but as not all hatchery fish are marked, wild fish could also encompass unmarked hatchery fish. DNA is extracted for detection of DNA viruses, bacteria and parasites from the same tissues from which we extract RNA to target RNA viruses. Nucleic-acid extractions on the aquaculture audit samples (8 tissues – gill, atrium, ventricle, liver, pyloric caeca, spleen, head kidney and posterior kidney) were as previously described (89). For the wild samples, homogenization using Tri-reagent™ was performed in a Mixer Mill (Qiagen, Maryland) on each tissue independently (5 tissues-gill, liver, heart, head kidney and brain). Tri-reagent™ homogenates were organically separated using bromochloropropane, with the RNA-containing aqueous layer removed for RNA extraction and the lower DNA-containing organic layer separated from the organics using a TNES-Urea Buffer (90).

For the DNA extractions, a pool of 250μl (5 tissues contributing 50 μl each) from each of the tissue TNES aqueous layers were processed for DNA using the BioSprint96 DNA Blood kit (Qiagen, Maryland) and the BioSprint96 instrument (Qiagen, Maryland) based on the manufacturer’s instructions. DNA was quantified using spectrophotometer readings performed on the Infinite M200Pro spectrophotometer (Tecan Group Ltd., Switzerland) and normalized to 62.5ng/μl using the Freedom Evo (Tecan Group Ltd., Switzerland) liquid handling unit, based on manufacturer’s instructions.

Similarly, a pool of 100μl (5 tissues contributing 20ul each) of the aqueous layer was processed for RNA using the Magmax™-96 for Microarrays RNA kit (Ambion Inc, Austin, TX, USA) with a Biomek NXPTM (Beckman-Coulter, Mississauga, ON, Canada) automated liquid-handling instrument, both based on manufacturer’s instructions. The quantity of RNA was analysed using spectrophotometer readings and normalized to 62.5ng/μl with a Biomek NXP (Beckman-Coulter, Mississauga, ON, Canada) automated liquid-handling instrument, based on manufacturer’s instructions. Mixed tissue RNA (1μg) was reverse transcribed into cDNA using the superscript VILO master mix kit (Invitrogen, Carlsbad, CA) following the manufacturer’s instructions.

### Metatranscriptomic sequencing

Samples which were VDD positive but were not positive for any known viruses based on RT-PCR screening (as described in (24)) were prepared for metatranscriptomic sequencing of RNA as previously described (12). RNA-seq libraries were prepared using the ScriptSeq Complete Epidemiology NGS library kit (Illumina, San Diego, CA), barcoded, and 4 samples were combined into RNA-seq runs on the Illumina MiSeq platform (Illumina, San Diego, CA).

Sequences were processed as previously described (12). In brief, host reads were removed by mapping to the Atlantic-salmon genome and unmapped reads were assembled *de novo* using SPAdes (v3.9.1) genome assembler (91); putative viral sequences were identified based on a similarity search. Assembled sequences are available on Genbank (Genbank accession numbers MN995807-MN995818). In some cases *de novo* assemblies were extended or scaffolded within Geneious. Assemblies were verified by re-aligning reads to the final scaffold to ensure there was continuous coverage of scaffold.

### Phylogenetic Analysis

To infer the phylogenetic relationship of the viruses sequenced from the salmon, the putative viral sequences were compared to sequences obtained from GenBank which showed similarity to our emerging salmon virus, or using sequences from the same viral family. False positives such as sequences with high similarity to plant viruses or host derived sequences which show similarity to viral sequences (endogenous viral elements) were not included. The evolutionary histories of the viruses were based on the predicted RdRp amino acid sequences; complete coding sequences for ASCV and CTV-2, and partial sequences for the incomplete viral genomes. Amino acid alignments were generated by MAFFT using the E-INS-i algorithm (92) and phylogenetic trees were generated with MrBayes (93). Trees were midpoint rooted for clarity only.

### RT-PCR

Assembled viral sequences from the appropriate sample were imported into Primer ExpressTM v3.0.1 software (Thermo Fisher Scientific, Waltham, MA) where qPCR Taqman assays were designed using default parameters. High-throughput RT-PCR using these assays was carried out as previously described (12, 94). For each assay, a theoretical limit of detection was applied which results in removing positive detections with a very low load. The limit of detection is set to the concentration of the analyte in the sample matrix that would be detected with high statistical certainty (95% of the time), as previously described (94).

### Sampling / Sample selection

A statistical model based on the hypergeometric distribution (95) was developed to assess the probability p that a particular novel virus would have been detected via sequencing. The hypergeometric distribution determines the chance that in a pond of m infected and n non-infected salmon (total m+n), x infected salmon are found when k salmon are pulled randomly from the pond. The probability p is equivalent to the chance that at least one infected salmon is found which can be derived as 1 minus the probability that no infected salmon is found in the k randomly pulled salmon. Figure 1A shows probability curves for different virus prevalence as a function of the number k of pulled salmon. All curves are based on a total of m+n=665 samples as an example, which is the number of Atlantic salmon in the farm audit study. For the curve with a virus prevalence of 10%, parameters m=67 and n=598 are used for the calculation with the hypergeometric distribution.

## Acknowledgements

We thank DFO Aquaculture Management Division, Salmon Enhancement Program, Environmental Watch Program, High Seas Program, Strait of Georgia salmon Program, PARR Program and the Hakai Institute Juvenile Salmon Program for provision of samples, and all of the vessel and field crews who collected the thousands of samples. This research was supported by funding for the Strategic Salmon Health Initiative (SSHI), which is part of the Salish Sea Marine Survival Project. The SSHI seeks to discover microbes present in salmon in British Columbia that may be undermining the productivity of Pacific salmon, and is funded by the Pacific Salmon Foundation (PSF), Genome British Columbia, and Fisheries and Oceans Canada through grants to KMS and CAS. GJM and EDC were supported by grants to KMS and CAS from MITACS and PSF, and computational resources provided by the Gordon and Betty Moore Foundation through a grant to CAS.

