## Supplementary materials for "Emerging viruses in British Columbia salmon discovered via a viral immune response biomarker panel and metatranscriptomic sequencing"

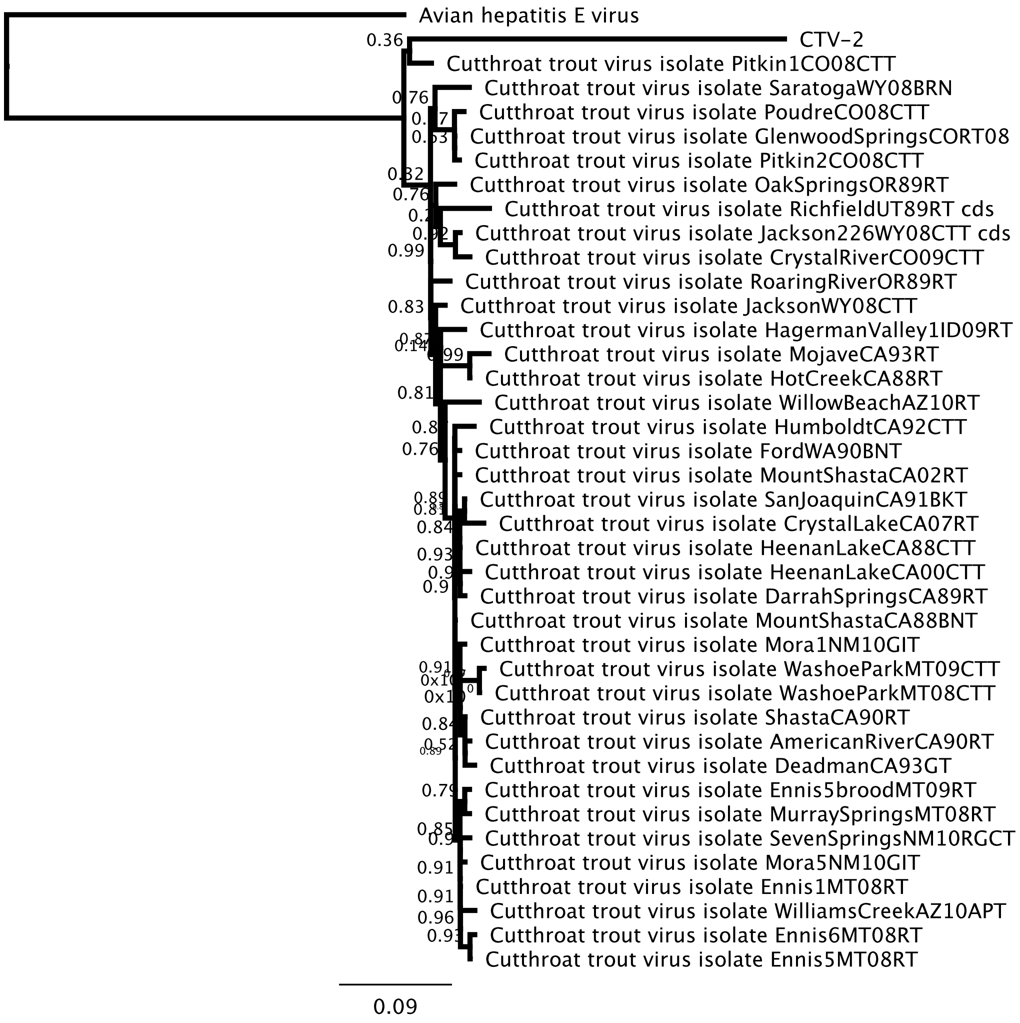


**Supplementary figure 1** Phylogenetic relationship of Cutthroat trout virus based on a portion of the helicase nucleotide sequence.

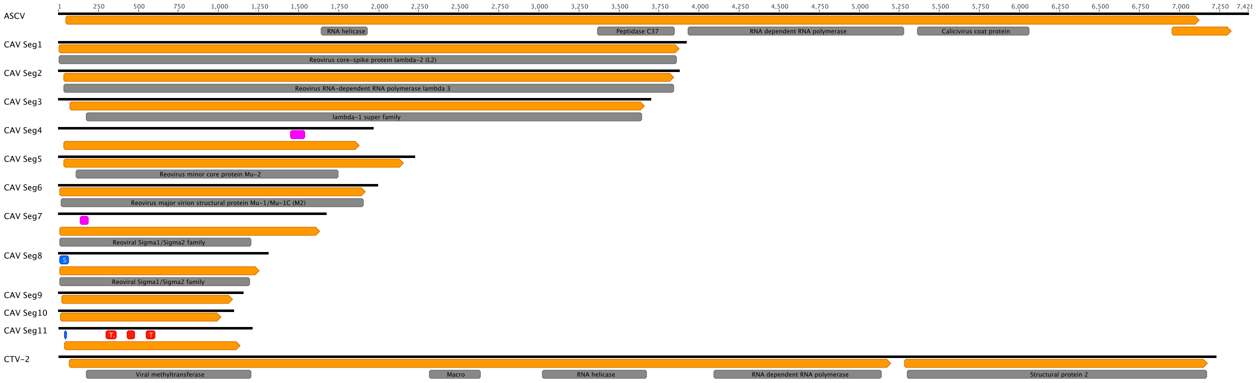


**Supplementary figure 2** Overview of the genome structure of the emerging viruses for which we sequenced coding complete genomes; Atlantic Salmon Calicivirus (ASCV), Chinook Aquareovirus (CAV) segments 1-11, and Cut-throat trout virus-2 (CTV-2). Black lines represent the genome, orange boxes show the predicted ORFs with conserved protein domains shown in grey. Blue boxes indicate the detection of a leader protein, pink boxes show predicted coiled-coil domains, and red boxes show predicted transmembrane domains.

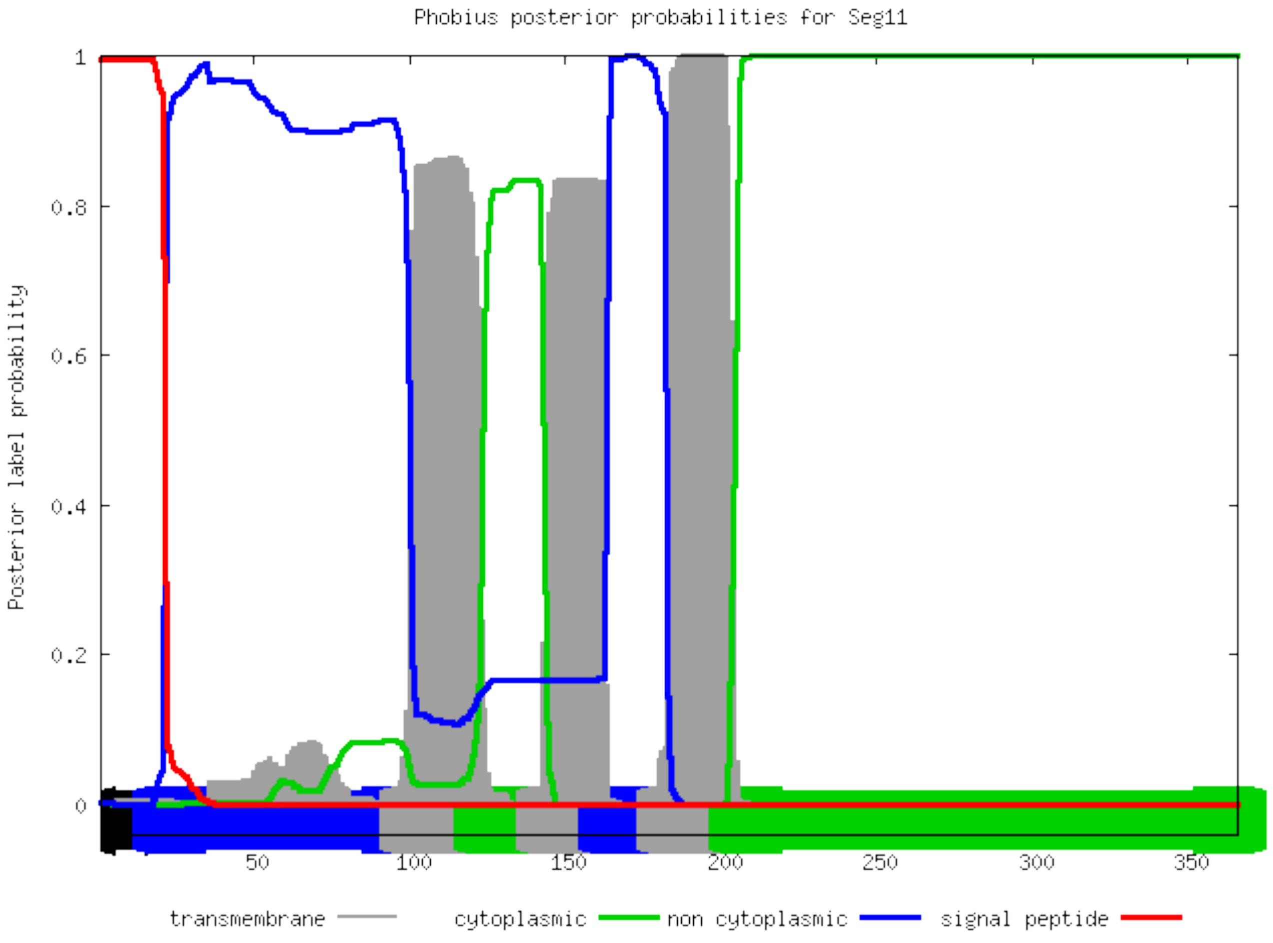


**Supplementary figure 3** Phobius output for the predicted protein encoded by Segment 11 of Chinook Aquareovirus (CAV)

| Segment | GC% | Length (nt) | Length (aa) | Homology |
|---|---|---|---|---|
| Seg-1 | 50.4086 | 3916 | 1292 | GCRV104 VP1 Core Turret |
| Seg-2 | 48.7232 | 3877 | 1272 | GCRV104 VP2 Core RdRp |
| Seg-3 | 50.1757 | 3699 | 1198 | GCRV104 VP3 Core shell |
| Seg-4 | 53.1091 | 1962 | 617 | GCRV104 VP66 NS factory |
| Seg-5 | 51.0791 | 2224 | 708 | GCRV104 VP5 Core NTPase |
| Seg-6 | 48.8956 | 1992 | 638 | GCRV104 VP4 Outer shell |
| Seg-7 | 48.5012 | 1668 | 543 | No homology – 'putative fiber protein' |
| Seg-8 | 50.3817 | 1310 | 417 | GCRV104 VP6 Core clamp |
| Seg-9 | 50.6957 | 1150 | 358 | GCRV RNAB |
| Seg-10 | 51.8282 | 1094 | 336 | GCRV104 outer clamp |
| Seg-11 | 50.6633 | 1206 | 366 | No homology – 'putative NS other' |

**Supplementary  Table 1** Chinook aquareovirus segments and homology to Hubei Grass Carp Reovirus

| Sequence Name | Predicted Protein function/ structure | Minimum (nt) | Maximum (nt) |
|---|---|---|---|
| ASCV | RNA helicase | 1641 | 1939 |
| ASCV | Peptidase C37 | 3366 | 3853 |
| ASCV | Calicivirus coat protein | 5363 | 6069 |
| ASCV | RNA dependent RNA polymerase | 3933 | 5285 |
| CAV Seg1 | Reovirus core-spike protein lambda-2 (L2) | 7 | 3869 |
| CAV Seg2 | Reovirus RNA-dependent RNA polymerase lambda 3 | 34 | 3848 |
| CAV Seg3 | lambda-1 super family | 174 | 3650 |
| CAV Seg4 | Coiled-coil | 1448 | 1548 |
| CAV Seg5 | Reovirus minor core protein Mu-2 | 112 | 1753 |
| CAV Seg6 | Reovirus major virion structural protein Mu-1/Mu-1C (M2) | 16 | 1912 |
| CAV Seg7 | Coiled-coil | 138 | 196 |
| CAV Seg7 | Reoviral Sigma1/Sigma2 family | 11 | 1209 |
| CAV Seg8 | Signal peptide | 8 | 71 |
| CAV Seg8 | Reoviral Sigma1/Sigma2 family | 8 | 1202 |
| CAV Seg11 | Signal peptide | 41 | 58 |
| CAV Seg11 | Transmembrane domain | 431 | 486 |
| CAV Seg11 | Transmembrane domain | 548 | 614 |
| CAV Seg11 | Transmembrane domain | 300 | 369 |
| CTV-2 | Macro | 2315 | 2643 |
| CTV-2 | RNA helicase | 3021 | 3679 |
| CTV-2 | Viral methyltransferase | 174 | 1212 |
| CTV-2 | RNA dependent RNA polymerase | 4094 | 5144 |
| CTV-2 | Structural protein 2 | 5301 | 7177 |

**Supplementary table 2** Predicted protein functions or structure for the coding complete genomes of ASCV-BC, CAV and CTV-2
